# CDK8/19 Inhibition Attenuates G1 Arrest Induced by BCR-ABL Antagonists and Accelerates Death of Chronic Myelogenous Leukemia Cells

**DOI:** 10.1101/2023.09.25.559286

**Authors:** Alvina I. Khamidullina, Margarita A. Yastrebova, Alexandra V. Bruter, Julia V. Nuzhina, Nadezhda E. Vorobyeva, Anastasia M. Khrustaleva, Ekaterina A. Varlamova, Alexander V. Tyakht, Yaroslav E. Abramenko, Ekaterina S. Ivanova, Maria A. Zamkova, Jing Li, Chang-Uk Lim, Mengqian Chen, Eugenia V. Broude, Igor B. Roninson, Alexander A. Shtil, Victor V. Tatarskiy

## Abstract

Imatinib mesylate (IM) and other BCR-ABL tyrosine kinase inhibitors (BCR-ABLi) are the mainstay of chronic myelogenous leukemia (CML) treatment. However, activation of circumventing signaling pathways and quiescence may limit BCR-ABLi efficacy. CDK8/19 Mediator kinases have been implicated in the emergence of non-genetic drug resistance. Dissecting the effects of pharmacological CDK8/19 inhibition on CML survival in response to BCR-ABLi, we found that a selective, non-toxic CDK8/19 inhibitor (CDK8/19i) Senexin B (SenB) and other CDK8/19i sensitized K562 cells to different BCR-ABLi via attenuation of cell cycle arrest. In particular, SenB prevented IM-induced upregulation of genes that negatively regulate cell cycle progression. SenB also antagonized IM-activated p27^Kip1^ elevation thereby diminishing the population of G1-arrested cells. After transient G1 arrest, cells treated with IM+SenB re-entered the S phase, where they were halted and underwent replicative stress. Consequently, the combination of IM and SenB intensified apoptotic cell death, measured by activation of caspase 9 and 3, subsequent cleavage of poly(ADPriboso)polymerase 1, positive Annexin V staining and increase of subG1 fraction. In contrast, IM-treated BCR-ABL-positive KU812 CML cells, which did not induce p27^Kip1^, readily died regardless of SenB treatment. Thus, CDK8/19i prevent the quiescence-mediated escape from BCR-ABLi-induced apoptosis, suggesting a strategy for avoiding the CML relapse.

## Introduction

The main genetic marker of chronic myelogenous leukemia (CML) is the Philadelphia chromosome generated by translocation t(9;22)(q34;q11). This translocation yields the oncogenic BCR-ABL fusion protein (1,2). The BCR-ABL chimeric tyrosine kinase activates a number of downstream pathways that drive the pathogenesis of CML as well as other tumor types including a subset of acute lymphoblastic leukemia, acute myelogenous leukemia, and mixed-phenotype acute leukemias (3,4).

Introduction of the tyrosine kinase inhibitor imatinib mesylate (IM, Gleevec) has drastically improved CML treatment outcomes (5). However, a significant proportion of patients eventually develop resistance to BCR-ABL inhibitors (BCR-ABLi). Such resistance was attributed to mutations that alter the drug-target interaction and/or to the involvement of BCR-ABL-independent signaling pathways (6). Second generation BCR-ABLi such as nilotinib, dasatinib and bosutinib, are used in the treatment of CML patients with certain mutations in the *ABL* gene, but can be inefficient in cells carrying the gatekeeper T315I mutation (7). While the third generation BCR-ABLi ponatinib, asciminib and PF-114 (vamotinib) are effective against specific *BCR*LJ*ABL* mutants including T315I, these drugs are limited in their ability to circumvent BCR-ABL-independent drug resistance (8).

BCR-ABL-independent resistance is mediated by signaling pathways that involve STAT3 (9,10), MAPK/ERK (11), β-catenin (12), as well as activation of autophagy (13). Dormant leukemia stem cells (LSC) capable of prolonged persistence as IM resistant cells (14) are especially refractory to BCR-ABLi. In CML, quiescence of LSCs is a mechanism that can prevent or delay the achievement of full clinical remission (15,16). Moreover, quiescent CML cells may re-enter the cell cycle, leading to a relapse after BCR-ABL targeted therapy (17). Cell cycle arrest and quiescence are regulated by cyclin dependent kinase inhibitor (CKI) proteins of CIP/KIP (p27^Kip1^, p57^Kip2^, and p21^Cip1^) and INK4 (p18^INC4c^ and others) families (18,19). BCR-ABL abrogates the p27^Kip1^ function (20–23), and inhibition of BCR-ABL by IM induces p57^Kip2^ followed by overexpression and stabilization of p27^Kip1^ (24). These data indicate that CML resistance to BCR-ABLi associated with cell cycle perturbations can be regulated by transcription and epigenetic regulation of CKI. Indeed, pharmacological intervention into the mechanisms of epigenetic modulation has been shown to be beneficial for the exit from quiescence, therefore increasing tumor cell death (25).

The cyclin dependent kinase 8 (CDK8) or its paralog CDK19, together with cyclin C, MED12, and MED13, form a kinase module associated with the multiprotein Mediator complex (26). This module regulates gene expression both positively and negatively by tuning the transcriptional machinery via the Mediator and transcription factor function at enhancers and promoters (27–29). The CDK8/19 module acts as a cofactor or modifier of different cancer-relevant transcription factors and coordinates the response to exogenous stimuli by reprogramming gene expression for optimal cell adaptation (reviewed in (28)). While CDK8/19 potentiate the induction of transcription by several signals, these kinases also inhibit Mediator-dependent transcription of super-enhancer associated genes which underlies their role in a subset of leukemias (30). CDK8/19 can modulate the transcription factors β-catenin (31), STAT1/3/5a (32,33), c-Myc (34), SMAD1/3 (35), NF-κB (36), and others. Importantly, CDK8 depletion does not affect the viability of adult cells or organisms (37,38), making these enzymes attractive drug targets.

The role of CDK8/19 in non-genetically acquired antitumor drug resistance has been demonstrated by the ability of selective small-molecule CDK8/19 inhibitors (CDK8/19i) to sensitize tumor cells or to prevent the emergence of resistance to chemotherapeutics including inhibitors of EGFR, HER2 and MEK (39–42). Furthermore, CDK8/19i potentiated growth inhibitory effects of the estrogen antagonist fulvestrant in breast cancer cells (43), reversed castration resistance of advanced prostate cancers (44), stimulated tumor surveillance by NK cells (45) and enhanced the antitumor effects of CAR-T cells (46). Thus, CDK8/19i are considered as antitumor drug candidates in combination regimens (reviewed in (47)). Several CDK8/19i have reached clinical trials in patients with solid tumors and leukemias (clinicaltrials.gov NCT03065010, NCT04021368, NCT05052255, NCT05300438).

Аn emerging role of CDK8/19 relates to S phase control in response to DNA replication defects defined as replication stress (RS). CDK8/19 module deficiency was reported to promote a premature entry into the S phase (48) and R-loop formation (49). However, another group reported that the depletion of cyclin C or CDK8 decreased the collisions between the transcription and replication machineries caused by inhibitors of RS response (50). Activities of CDK8/19 in RS response might be context-dependent, an implicit feature of these kinases (36).

In the present study, we discovered the ability of CDK8/19i to sensitize CML cells to BCR-ABLi via prevention of BCR-ABLi-induced cell cycle arrest. Cells treated with the combination of CDK8/19i and BCR-ABLi enter replication, leading to an increase of RS markers and accelerating the onset and the rate of apoptosis. These results suggest that inhibition of CDK8/19 may prevent quiescence-mediated resistance to BCR-ABLi.

## Materials and Methods Reagents

All reagents were from Sigma-Aldrich, Burlington, MA unless specified otherwise. IM (Gleevec®) was purchased from Novartis, Basel, Switzerland. Dasatinib and nilotinib were from Selleck Chemicals, Houston, TX. PF-114 (51), recently named vamotinib, was a gift of Dr. G. Chilov (Valenta Pharm, Moscow). CDK8/19i Senexin B (SenB) and SNX631 were from Senex Biotechnology, Columbia, SC. Drugs were dissolved in dimethyl sulfoxide (DMSO) as 10 mM stock solution and stored at -20°C. Dilutions in the culture medium were prepared immediately before experiments.

### Cell lines and culture conditions

Human CML cell lines K562 (Russian Collection of Cell Cultures, Saint-Petersburg, Russia) and KU812 (CRL-2099-ATCC, Manassas, VA) were propagated in RPMI-1640 (PanEco, Moscow, Russia) with 10% fetal bovine serum (Biosera, Cholet, France), 2 mM *L*- glutamine, 100 U/mL penicillin and 100 μg/mL streptomycin (PanEco) at 37°C and 5% CO_2_ in humidified atmosphere. Cells (2×10^5^ cells/mL) in the logarithmic phase of growth were plated into 60 mm Petri dishes (SPL, Korea) and treated with CDK8/19i or BCR-ABLi alone (concentrations and time of exposure are indicated in respective experiments) or with the combination of the two agents (CDK8/19i added 1 h apart from BCR-ABLi). Control wells contained 0.02% DMSO (vehicle).

### Flow cytometry

Cell cycle distribution was analyzed as described (52). After the completion of treatments cells were washed with cold saline, pelleted and lysed in a buffer containing 50 μg/mL propidium iodide (PI), 100 μg/mL RNAse A, 0.1% sodium citrate, 0.3% NP-40 (VWR Life Science, Radnor, PA) for 30 min in the dark. Apoptosis was analyzed with Apoptotic, Necrotic & Healthy Cells Quantitation Kit Plus (Biotium, Fremont, CA) in accordance with the manufacturer’s recommendations. The MitoTracker® Red CMXRos (Invitrogen, Carlsbad, CA) was used to evaluate the mitochondrial membrane potential. Briefly, cells were stained with 500 nM MitoTracker solution in the medium for 40 min at 37[, 5% CO_2_ in the dark, washed with warm medium and analyzed by flow cytometry. Live/dead cells were determined by PI Nucleic Acid Stain (Thermo FS, Waltham, MA). Survival fraction was calculated as the percentage of PI negative cells after normalizing to the total cell count. To assess proliferation, cells were labeled with 10 μM 5-ethynyl-2’- deoxyuridine (EdU) for 2 h and analyzed using ClickTech EdU Cell Proliferation Kit 488 (Carl Roth, Karlsruhe, Germany) according to the manufacturer’s recommendations. Fluorescence was measured on a Cytoflex flow cytometer 26 (Beckman Coulter, Indianapolis, IN) in FITC, APC, PE-A and PerCP-A channels. At least 10,000 events were collected per each sample. Data were analyzed using CytExpert Software (Beckman Coulter).

### RNA sequencing (RNASeq)

#### Preparation of cDNA libraries

K562 cells (2×10^5^ cells/mL) were treated with the vehicle (0.02% DMSO), 1 μM SenB, 1 μM IM or their combination (two replicates per each treatment) for 8 h. The vehicle and SenB were added 1 h prior to TKI. Total RNA was extracted with TRI reagent. 4 μg RNA was used to isolate poly(A)-enriched RNA with NEBNext® Poly(A) mRNA Magnetic Isolation Module (NE Biolabs, Ipswich, MA) that was used to prepare RNASeq libraries with NEBNext® Ultra™ II Directional RNA Library Prep Kit for Illumina (NE Biolabs). Actinomycin D (100 ng/mL) was used for first strand cDNA synthesis; cDNA libraries ligated with the Illumina suitable adaptor sequences were generated and amplified with Q5 DNA polymerase (NE Biolabs). After purification from dimers by size selection in the agarose gel the libraries were sequenced on a NovaSeq 6000 (Illumina, San Diego, CA).

#### Analysis of RNASeq data

Table S1 lists the references to the software and algorithms mentioned below. Quality of short reads was checked using the FastQC software. Reads were trimmed (i.e., sequencing adaptors removed) using cutadapt, reads with a Phred quality score[<[20 and read length[<[30 bp were removed using the sickle. Mapping of trimmed reads to human genome assembly GRCh37 (hg19) and calculation of per-gene read counts were performed using STAR.

Statistical analysis was conducted using the edgeR package. Only genes with counts greater than one per million (cpm>1) in both samples were included. Read counts were normalized using the trimmed mean of values method implemented in edgeR to account for differences in the library size. General linear models and the likelihood ratio test were used to identify differentially expressed genes (DEGs). The Benjamini-Hochberg false discovery rate (FDR) correction was applied to the test results (alpha=0.05).

Over-representation analysis (ORA) using both Gene Ontology (GO) enrichment analysis of DEGs and Reactome Pathways Database were conducted using the WebGestaltR package. The gene set enrichment analysis (GSEA) for different comparisons was conducted using the fgsea package with the specific gene sets downloaded from the Human Molecular Signatures Database (MsigDB). The ggplot2 and clusterProfiler R libraries were used for data visualization.

All raw RNASeq data have been uploaded to Sequence Read Archive (SRA, SUB13787508) and BioProject (PRJNA1008677). Detailed information about RNASeq samples is listed in Table S2.

#### Lentiviral transduction

To obtain pCW-p27 lentiviral plasmid carrying *CDKN1B*/p27^Kip1^ open reading frame (ORF; RefSeq NM_004064.5), total RNA was isolated from IM-treated K562 cells. Primers CDKN1B-for 5’-attagctagcATGTCAAACGTGCGAGTGTCTAA-3’ and CDKN1B-rev 5’-taatggatccTTACGTTTGACGTCTTCTGAGGC-3’ (Evrogen, Moscow, Russia) containing NheI and BamHI restriction sites were used for amplification of the complementary DNA. The ORF was then cloned into the pCW vector replacing Cas9 in the pCW-Cas9 plasmid (https://www.addgene.org/50661/). The K562p27tet-on subline with doxycycline inducible *CDKN1B*/p27^Kip1^ overexpression was obtained by lentiviral transduction. The virus was concentrated by ultracentrifugation (120 000 g) for 2 h at 4°C. Polybrene (20 µg/mL) was added, and the supernatant was mixed with K562 cells in the fresh medium (1:1 v/v). Selection was performed with 2 µg/mL puromycin. The exogenous p27^Kip1^ was induced by 1 µg/mL doxycycline.

#### Immunoblotting

K562 and KU812 cells (2×10^5^ cells/mL) were treated with 0.02% DMSO or drugs, harvested and lysed for 30 min on ice in a buffer containing 50 mM Tris-HCl pH 8.0, 150 mM NaCl, 0.1% sodium dodecyl sulfate, 1% NP-40, 2 mM phenylmethylsulfonyl fluoride (VWR Life Science) and the protease inhibitor cocktail. Protein concentrations were determined by the Bradford method. Lysates were separated by SDS-PAGE (30-50 μg total protein per lane) and transferred onto a 0.2 μm nitrocellulose membrane (Bio-Rad, Hercules, CA). Membranes were blocked with 5% skimmed milk for 30 min and treated with primary antibodies (listed in Table S3) diluted in Tris-borate saline/Tween 20 (TBST)/1% bovine serum albumin (PanEco) overnight at 4°C. Then membranes were washed with TBST and incubated with a secondary antibody conjugated with horseradish peroxidase (Table S3) for 1 h at room temperature. Proteins were visualized with the Clarity Western ECL Substrate (Bio-Rad) using the iBright FL1500 Imaging System (Invitrogen).

#### Statistical analysis

Data are representative of at least three independent experiments. One-way or two-way analyses of variance (ANOVA) followed by Sidak’s post hoc test for multiple comparisons were used (GraphPad Prism 9; GraphPad Software, San Diego, CA). The P value <0.05 was taken as evidence of statistical significance.

## Results

### Senexin B increases IM-induced apoptosis in K562 cells

We tested whether a selective CDK8/19i SenB (43,53) affects CML cell viability in response to BCR-ABLi IM. Hereafter, cells were pre-treated with 1 μM SenB for 1 h followed by the addition of BCR-ABLi. Figure 1A, *left* shows that SenB sensitized K562 cells to IM. By 72 h of exposure the percentage of viable cells was larger in the IM-treated cohort than in the combination. The portion of propidium iodide (PI) positive (late apoptosis) cells greatly increased from 15.0±0.3% (IM alone) to 54.0±0.6% in the combination of IM and SenB (Figure 1A, *right*; p<0.0001). The time course showed that, already by 24 h, 1 μM SenB strongly increased the percentage of subG1 events (cells with fragmented DNA) after treatment with low concentrations of IM (Figure 1B, *left*; compare 11.6±2.2% in cells treated with 0.25 μM IM *vs* 24.6±1.4% after 0.25 μM IM and 1 μM SenB (p<0.0001)). Because the maximum difference between IM alone and IM+SenB was detected by 24 h, this time point was used in the next experiments. Potentiation of apoptosis was observed even with submicromolar concentrations of SenB (Figure 1B, *right*). SenB alone did not increase the percentage of Annexin V-positive cells but cooperates with IM in elevating this fraction (Figures 1C, *left*, S1). In contrast to K562 cells, the BCR-ABL positive KU812 CML cell line was intrinsically hypersensitive to IM; SenB had no significant effect on the already very high apoptotic fraction in cells treated with IM alone (Figures 1C, *right*, S1).

**Figure 1.**
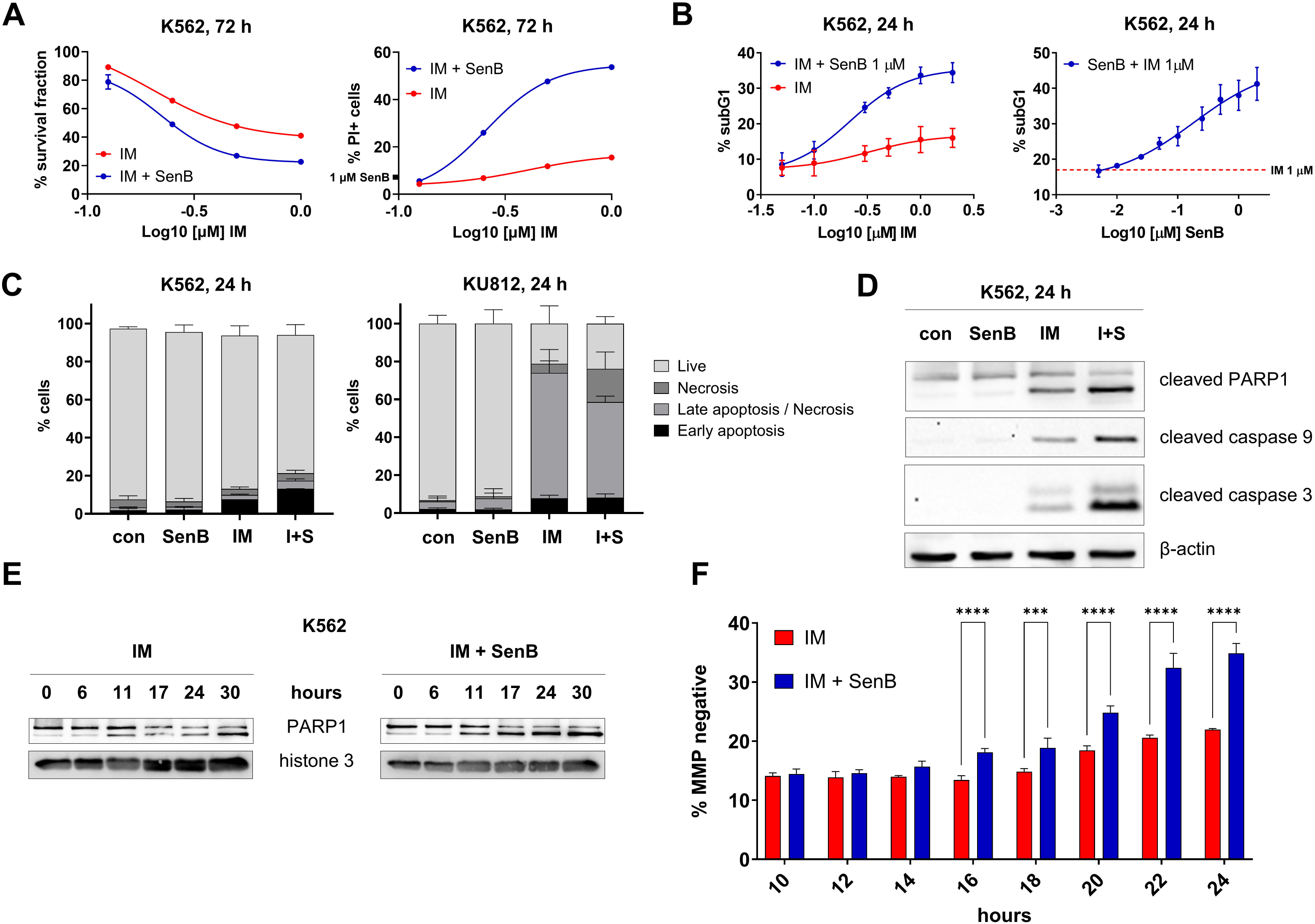
SenB cooperates with IM in inducing apoptosis in K562 cells. Cells were treated with SenB, IM (1 µM each) or their combination (I+S) for indicated time intervals. Flow cytometry assisted analysis showed **(A)** a decreased total cell number in I+S combination and a significant increase in PI+ cells compared to IM alone; **(B)** dependence of subG1 fraction on concentrations of SenB and IM; (С) increase of Annexin V-positive cells in I+S *vs* IM cohort in K562 cells (*left)* but not in KU812 cells (*right)*. Immunoblotting demonstrates **(D)** increased cleavage of PARP1, caspases 9 and -3 (apoptotic markers) in I+S *vs* IM alone. Β-actin, a loading control; (**E**) a faster onset of PARP1 cleavage in I+S combination *vs* IM alone. Histone 3, a loading control. **(F)** Time course of MMP changes determined by flow cytometry. Statistical analysis was performed using a two-way ANOVA. ***p<0.001, ****p<0.0001. Values are mean±SD, n=3.

Next, we found that the combined inhibition of CDK8/19 and BCR-ABL increased the proteolytic cleavage of poly(ADPriboso)polymerase 1 (PARP1) in K562 cells (Figure 1D). Furthermore, the time course of cell death markers (Figure 1E) showed that PARP1 cleavage was more readily inducible by the combination of IM and SenB than by IM only.

Finally, SenB significantly increased IM-induced decrease of the mitochondrial membrane potential (MMP) already by 16 h (Figure 1F), further indicating that SenB accelerated the onset and the rate of IM-induced apoptosis in K562 cells.

SenB potentiated cell death induced not only by IM but also by BCR-ABLi of different chemical classes such as nilotinib, dasatinib, PF-114/vamotinib (Figures 2A-B). Additionally, the interaction with IM was observed for a chemically unrelated CDK8/19i SNX631 (41,54), suggesting that the sensitization is a general effect of CDK8/19 inhibition (Figures 2C-D).

**Figure 2.**
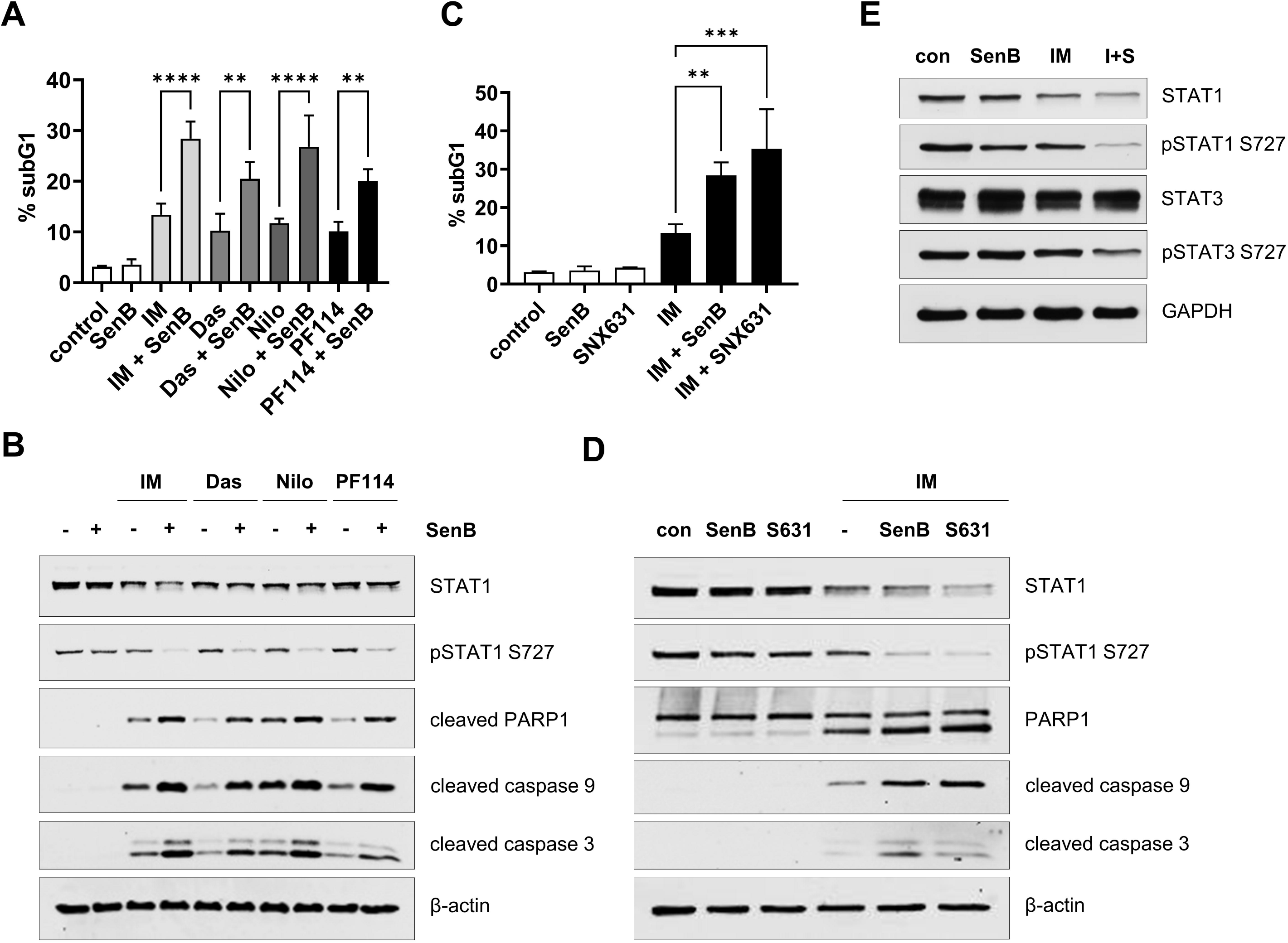
Combinations of CDK8/19i and BCR-ABLi cooperatively induce apoptosis and inhibit phosphorylated STAT1 and STAT3 S727 in K562 cells. K562 cells were treated with CDK8/19i and/or BCR-ABLi for 24 h. **(A)** Cell cycle distribution/PI staining followed by flow cytometry. Shown are the percentages of subG1 events in untreated cells (control) and cells treated with 1 µM IM, 1 nM dasatinib (das), 50 nM nilotinib (nilo) or 10 nM PF-114 alone or with 1 µM SenB. Note an increased subG1 fraction in combinations of all BCR-ABLi and SenB. **(B)** Immunoblotting showed that IM, das, nilo and PF-114, each in combination with SenB, inhibited STAT1 S727 phosphorylation and induced the cleavage of PARP1, caspases 9 and -3. SenB and SNX631 (S631) in combination with IM elevated **(C)** subG1 proportion and **(D)** apoptotic markers **(E)** SenB together with IM (I+S) decrease the amounts of pSTAT1 S727, STAT1 and pSTAT3 S727. GAPDH and β-actin are loading controls. Statistical analysis was performed using one- way ANOVA. **p<0.01, ***p<0.001, ****p<0.0001. Values are mean±SD, n=3.

The S727 site of STAT transcription factors is a known target of CDK8 that regulates STAT-mediated transcription (32,33,55). SenB or IM alone had no major effect on the amount of total STAT1 or pSTAT1 S727 in K562 cells, but both pSTAT1 S727 and total STAT1 were strongly decreased by the combination of IM and SenB (Figure 2E). Combined treatment with SenB and IM also reduced STAT3 S727 phosphorylation but not total STAT3.

### IM and SenB affect the expression of cell cycle related genes

RNASeq revealed changes in gene expression patterns after 8 h treatment with SenB or IM alone or in combination. We therefore analyzed the transcripts before the activation of cell death machinery to avoid apoptotic mRNA degradation. Comparisons between the effects of different treatments are shown in the volcano plots in Figures S2-S5. DEGs are listed in Tables S4-S5. IM induced downregulation of 2 100 genes and upregulation of 1 756 genes, whereas SenB downregulated 679 genes and upregulated 1 017 genes (Figures S2, S3). The combination of SenB and IM induced downregulation of 2 394 genes and upregulation of 2 073 genes (Figure S4). The addition of SenB changed the expression of 1 185 (downregulation) and 1 492 mRNAs (upregulation) compared to IM alone (Figure S5). Among the most affected genes (FC (fold change) > 1.5, FDR < 0.01) that significantly IM+SenB groups (Figure 3A). Interestingly, among the genes most strongly affected by IM, SenB most often enhanced their upregulation but counteracted their downregulation (Figure 3B). GSEA and ORA (Table S1) showed that many of the genes downregulated by IM were related to cell proliferation and interferon/STAT signaling (Figures 3C), coding for pro- proliferative proteins and cell cycle inhibitors. The addition of SenB evoked no major changes in the overall effects of IM on the hallmark pathways (Figure 3C), but many genes differentially affected by IM+SenB *vs* IM alone were associated with cell proliferation (Figure 3D, S6). Notably, 17.1% of all DEGs downregulated by the combination *vs* IM alone were genes related to negative regulation of cell cycle (according to GO terms, Table S6), while only 2.9% of genes related to cell cycle regulation were upregulated (Table S7). The top five cell cycle genes upregulated by IM (*VASH1*, *PCBP4*, *GPNMB*, *INHA* and *BTN2A2*) were all related to negative regulation (i.e., arrest) of the cell cycle, and were downregulated in IM+SenB treated cells (Figure S7). Therefore, a major effect of SenB in the combination was an attenuation of IM-induced negative cell cycle regulation.

**Figure 3.**
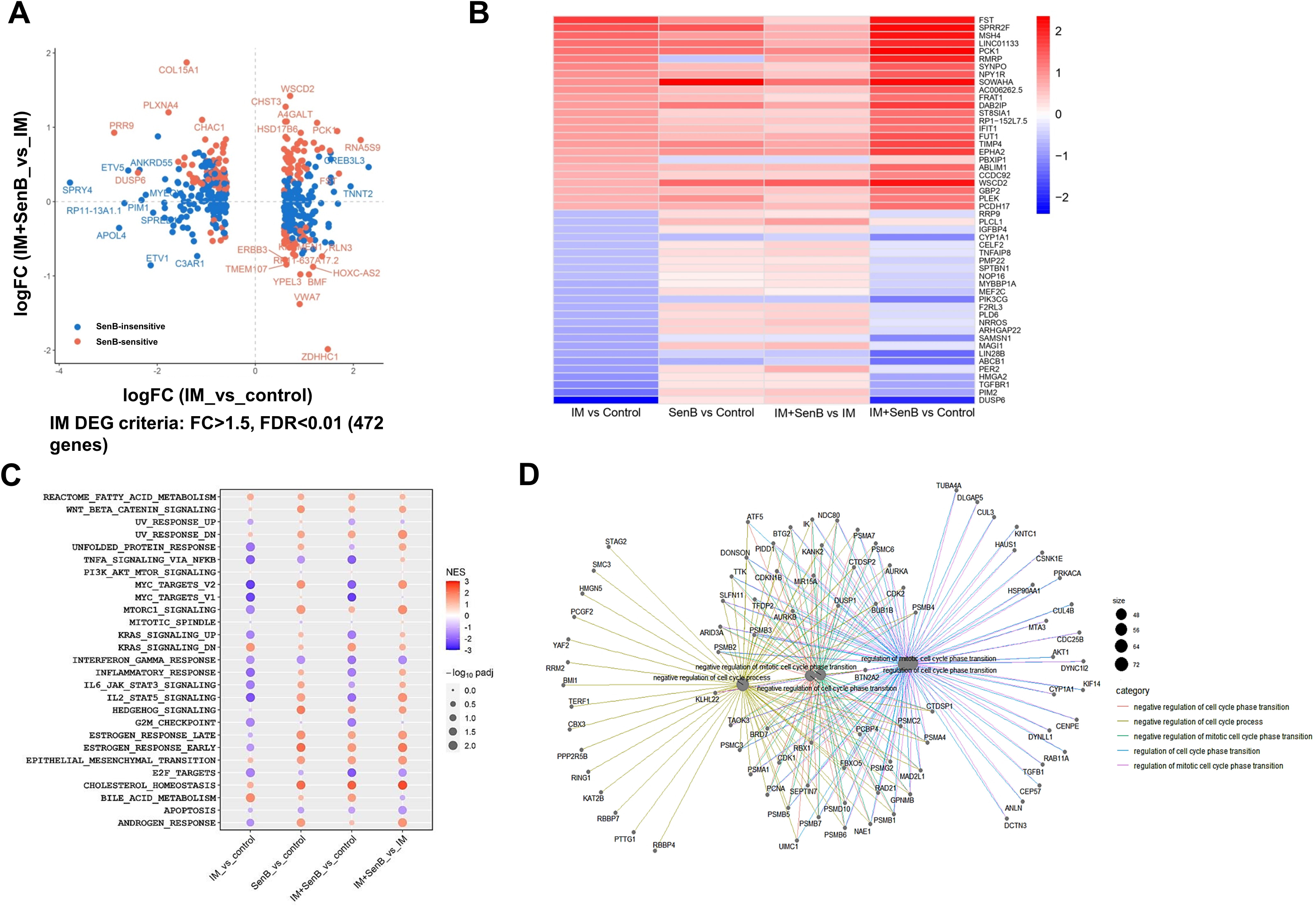
RNASeq analysis identifies the genes associated with the cell cycle. The K562 cells were treated for 8 h with SenB, IM (1 μM each) and their combination. **(A)** A dot plot comparing the effects of IM *vs* untreated control and IM+SenB *vs* IM on the differentially expressed genes (DEGs) affected by IM (FC (fold change) > 1.5, FDR (false discovery rate) < 0.01) shows both SenB-sensitive (red dots) and -insensitive genes (blue dots). 6.1% (163 genes) were differentially expressed in IM and IM+SenB groups. **(B)** The heatmap of the effects of different treatments on the genes most strongly upregulated or downregulated by IM demonstrates predominant upregulation of IM-induced genes and downregulation of IM-inhibited genes by the addition of SenB. **(C)** Analysis of hallmark pathways (GSEA). Combined treatment upregulated the majority of pathways compared to IM alone but did not prevent the IM-induced changes in the majority of pathways. (**D**) Netplot of genes downregulated in IM+SenB cohort *vs* IM alone. IM+SenB downregulates cell cycle related genes (17.1 % of all DEG downregulated by the combination *vs* IM alone).

### CDK8/19 inhibition abrogates IM-induced G1 arrest and elevates the markers of replication stress

According to the results of RNASeq analysis, the cell proliferation driver c-Myc was downregulated by IM, but upregulated by SenB; *MYC* expression in cells treated with IM + SenB remained the same as in untreated control (Figure 4A, *left*). In contrast, the cell cycle inhibitor *CDKN1B* (p27^Kip1^) was strongly upregulated by IM and downregulated by SenB, and the IM+SenB combination did not change *CDKN1B* expression (Figure 4A, *right*). p27^Kip1^, a major regulator of G1 checkpoint, is known to be involved in the response to IM (56,57). Figure 4B shows that IM treatment increased p27^Kip1^ protein in K562 cells in a dose-dependent manner whereas SenB strongly suppressed this effect. IM also upregulated two other CDK inhibitors, p57^Kip2^ and p18^INC4c^; the increase in p57^Kip2^ and p18^INC4c^ was barely sensitive to SenB (Figure 4C). Similarly, SenB only slightly reduced the effect of IM on the c-Myc protein (Figure 4C, *left*), similarly to the changes of c-Myc dependent transcripts (Figure 3C).

**Figure 4.**
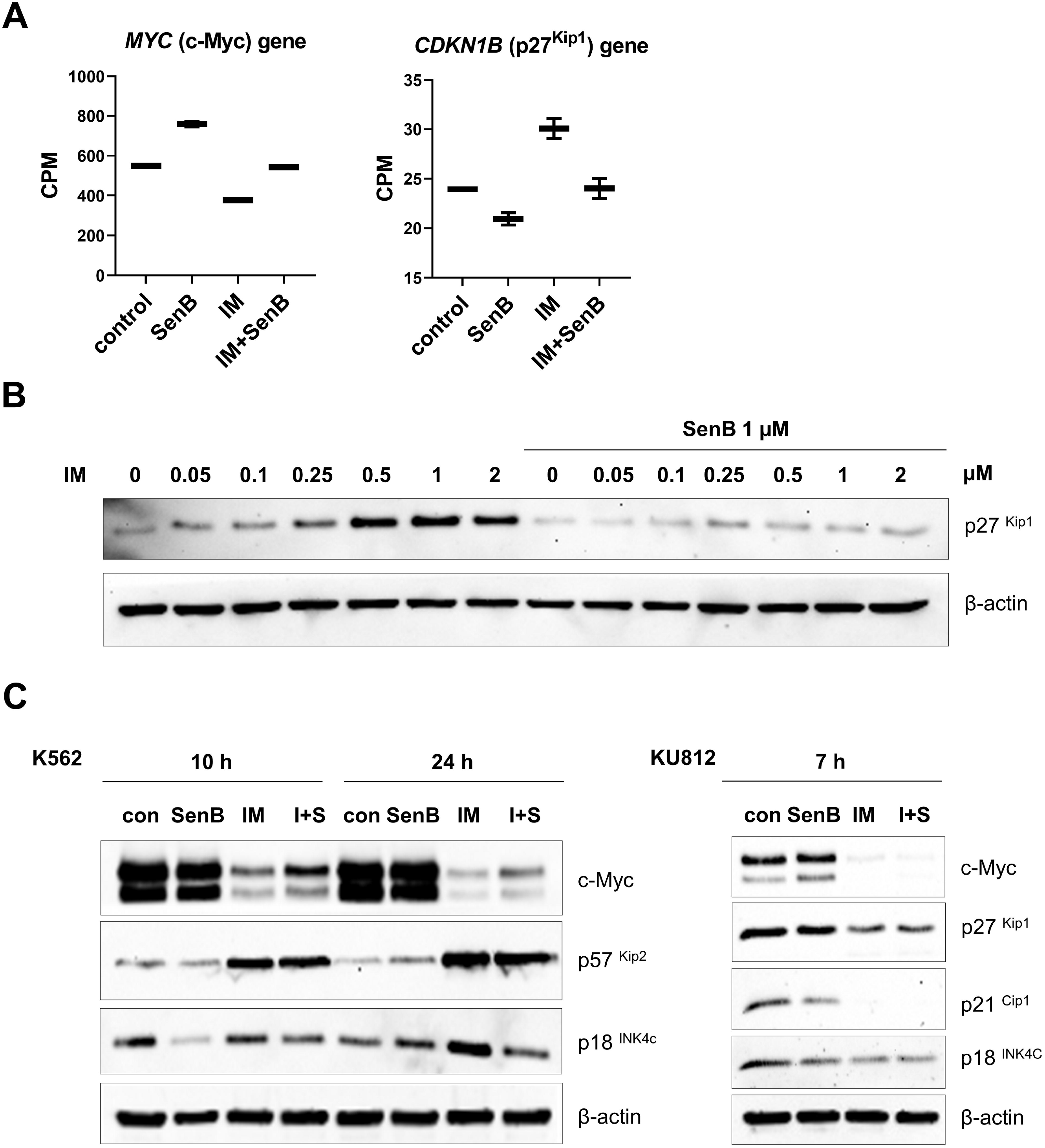
Combinations of IM and SenB affect cell cycle genes and proteins in K562 cells but not in KU812 counterparts. K562 and KU812 cells were treated as indicated. Transcripts were analyzed by RNASeq, and proteins by immunoblotting. (**A**) Cell proliferation and cell cycle markers analyzed by RNASeq. SenB in combination with IM counteracts *MYC* and *CDKN1B* (p27^Kip1^) upregulation by IM. The abundance of *MYC* and *CDKN1B* mRNAs is shown as normalized counts per million (CPM) reads. False discovery rate (FDR) < 0.001. (**B)** SenB prevents IM- induced p27^Kip1^ accumulation. (**C**) Immunoblotting of c-Myc, p57^Kip2^, p18^INC4c^ in K562 (*left*) induces accumulation of CKIs and downregulation of c-Myc in K562 cells, SenB in combination with IM (I+S) partially reverses this effect, but not in KU812 cells. β-actin is a loading control.

Given that IM potently upregulated p27^Kip1^ whereas SenB counteracted this effect, it is plausible that SenB-mediated prevention of p27^Kip1^ elevation by IM could contribute to the pro-apoptotic efficacy of the combination. In agreement with this hypothesis, a very different pattern was observed in IM-hypersensitive KU812 cells: p27^Kip1^, p18^INC4c^ and c-Myc were not induced by IM; instead, disappearance of c-Myc and p21^Cip1^ was detectable already by 7 h with IM; these effects were insensitive to SenB (Figure 4C, *right*).

To investigate whether p27^Kip1^ can mediate the sensitization of K562 cells to IM by SenB, we generated an inducible K562p27^tet-on^ derivative in which the exogenous *CDKN1B* is expressed under the control of doxycycline-inducible promoter. Before the addition of doxycycline, the percentage of K562p27^tet-on^ cells in G1 phase was 35.3±1.6%; by 24 h of doxycycline treatment, this value increased up to 64.8±0.7%, p<0.0001. SenB attenuated p27^Kip1^-induced G1 arrest in a dose dependent manner (Figure S8); 1 μM SenB decreased the G1 fraction to 52.4±2.2%. IM+doxycycline further elevated the G1 fraction, whereas SenB abrogated this increase (Figure 5A, *left*). This effect of SenB was paralleled by the increase of subG1 events (Figure 5A, *right*) and the fractions of cycling cells (S and G2/M phases) (Figure S9). As shown in Figure 5B, doxycycline-induced p27^Kip1^ partially prevented IM- induced apoptosis in K562p27^tet-on^ cells: 17.3±2.7% subG1 events in the ‘no doxycycline’ group *vs* 11.6±0.4% in the ‘doxycycline’ cohort, p<0.01. In contrast, the addition of SenB increased the percentage of the subG1 events to 26.4±3.4%, thereby overcoming the protective effect of p27^Kip1^ induction. SenB also partially decreased p27^Kip1^ levels with and without doxycycline (Figure 5C). Nevertheless, SenB did not fully abrogate the protective effect of p27^Kip1^ which can point to an alternative explanation that p27^Kip1^ is downregulated in cells entering replication. Hence, CDK8/19i acts through decreasing G1 arrest by alternative mechanisms.

**Figure 5.**
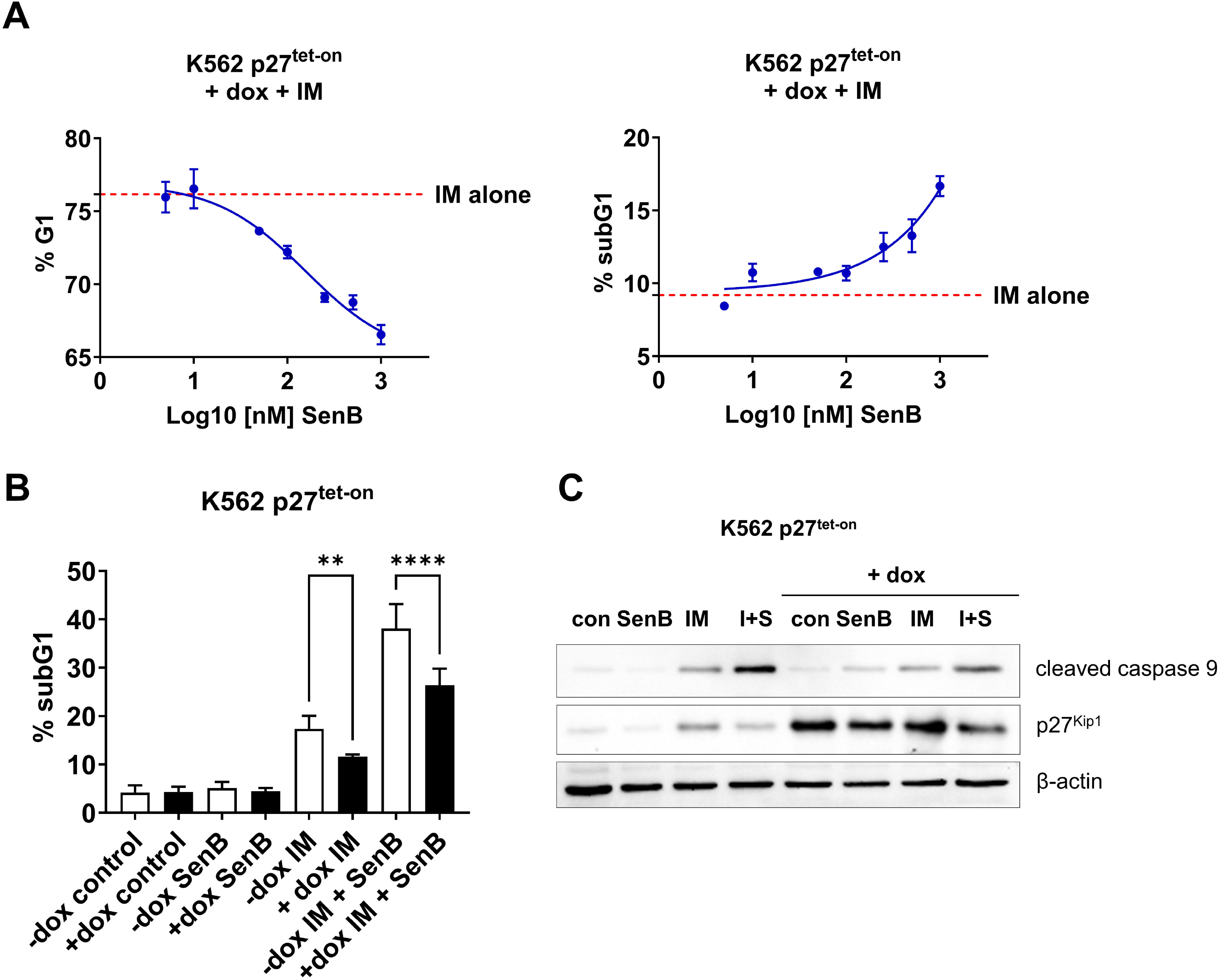
SenB counteracts the effects of the exogenous *CDKN1B* (p27^Kip1^) on cell cycle and IM survival. **(A)** K562p27^tet-on^ subline was treated with 1 µg/mL doxycycline (dox) and 1 µM IM for 24 h in the absence or presence of indicated concentrations of SenB followed by flow cytometry. SenB in a dose-dependent manner increases IM-induced apoptosis (subG1; *right*) and reduces G1 arrest *(left)*. **(B)** K562p27^tet-on^ cells were treated with IM, SenB (1 µM each) or their combination for 24 h in the absence or presence of vehicle (-dox) or 1 µg/mL doxycycline (+dox) followed by flow cytometry. Induction of p27^Kip1^ reduces IM-induced apoptosis (subG1 fraction). SenB partially alleviates the protective effect of p27^Kip1^ on IM-induced apoptosis. **(C)** Immunoblotting of cleaved caspase 9 and p27^Kip1^ in K562p27^tet-on^ cells treated with IM, SenB or their combination for 24 h in the absence or presence of vehicle (-dox) or 1 µg/mL doxycycline (+dox). Exogenous p27^Kip1^ overexpression decreases apoptosis marker after treatment with IM, addition of SenB (I+S) increases active caspase 9 level relative to IM alone. β-actin was used as a loading control. Statistical analysis was performed using one- way ANOVA. **p<0.01, ****p<0.0001. Values are mean±SD, n=3.

Next, we investigated the fate of cells that remained cycling upon treatment with IM or IM+SenB combination. As shown in Figure 6A (*top left*), by 10 h of treatment, K562 cells treated with IM also underwent G1 arrest similarly to IM+SenB treated counterparts. However, while G1 arrest by IM alone was maintained for the entire 32 h period of observation, this arrest was transient in cells treated with IM+SenB, and subsequently a portion of cells re-entered the cycle. The cell cycle re-entry in combination-treated cells was paralleled by an increased apoptosis rate than in cells treated with IM alone (Figure 6A, *bottom left*). In contrast to K562 cells, the highly IM sensitive KU812 cells did not undergo G1 arrest in response to IM (Figure 6A, *top right*) and readily died regardless of SenB treatment (Figure 6A, *bottom right*).

**Figure 6.**
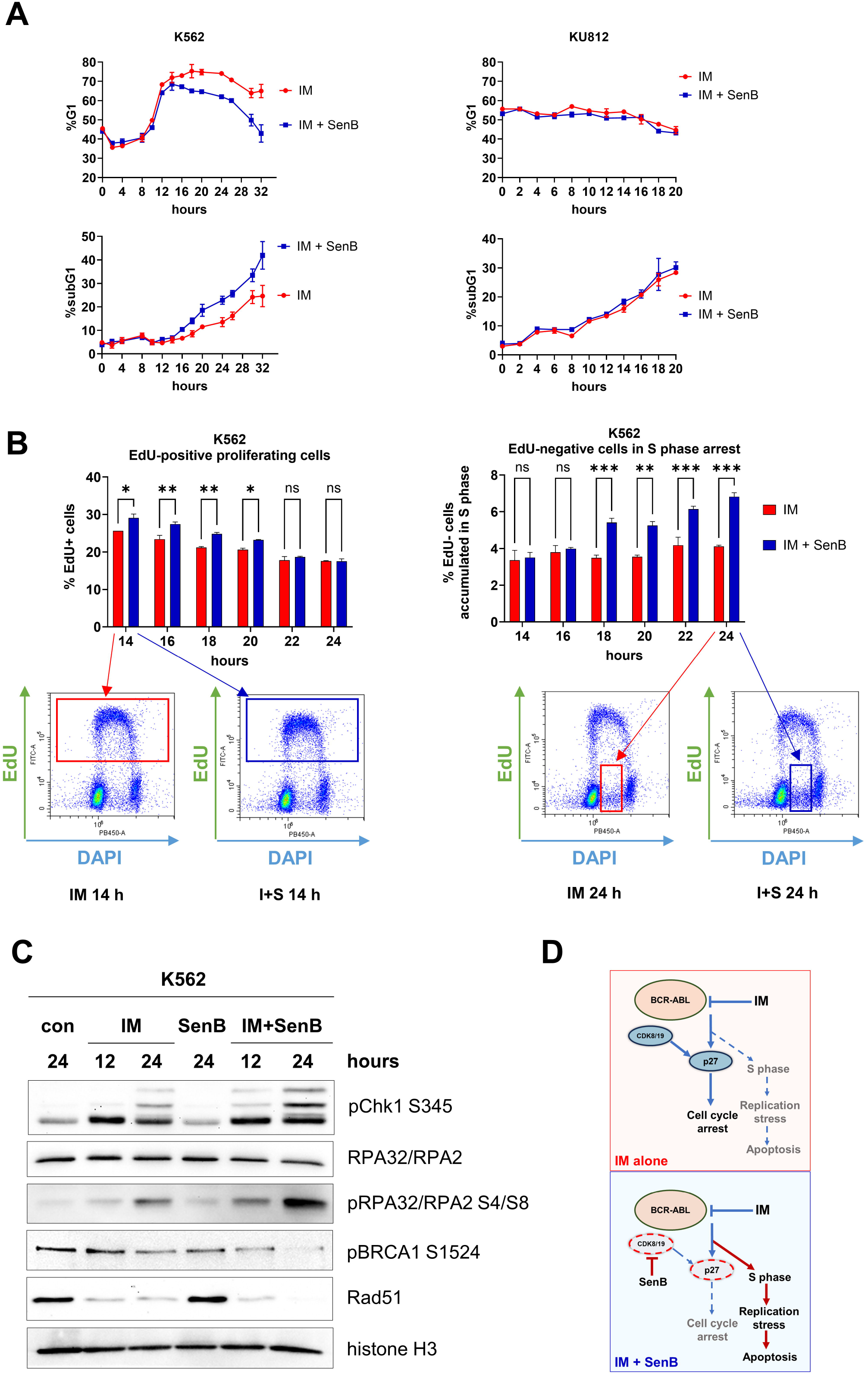
SenB alleviates IM-induced G1 arrest and increases replication stress markers. **(A)** Time course of G1 and subG1 fractions of K562 (*left*) and KU812 (*right*) cells treated with IM and SenB (1 µM each). Note that, in K562 cells, IM alone caused a sustained accumulation in G1 whereas in the IM+SenB group a portion of cells in G1 gradually the ‘IM alone’ cohort (*bottom left*). Since KU812 cells were not arrested by IM, the combination had no effect (*top & bottom right*). **(B)** Time course of EdU incorporation (2 h pulse) in K562 cells treated with IM and IM+SenB (I+S). Cells treated with IM+SenB re- started replication over 14-20 h (*left*), but at 18 h onward were arrested in the S phase (*right*). Shown is one representative experiment out of three biological replicates (*bottom*). **(C)** Immunoblotting of replication stress markers pChk1 S345, RPA32/RPA2, pRPA32/2 S4/8, Rad51, pBRCA1 S1354. Combination IM+SenB exacerbated the effects of IM on replication stress markers and inhibited the BRCA1-Rad51 DNA repair pathway. Histone 3 was a loading control. **(D)** Proposed mechanism of action of IM+SenB combination. Statistical analysis was performed using a two-way ANOVA. *p<0.05, **p<0.01, ***p<0.001; ns, not significant. Values are mean±SD, n=3.

We used EdU labeling of replicating cells to test whether K562 cells treated with IM or IM+SenB died after entering the S phase after G1 arrest or directly in G1. The fraction of EdU-positive cells was determined every 2 h over 12 h, starting at the 12 h point after the addition of drugs. As shown in Figure 6B, *left*, EdU-positive fraction increased in cells treated with IM+SenB relative to IM-treated cells at 14-20 h. After 20 h the amounts of EdU were similar in IM and IM+SenB groups, but the percentage of EdU-negative cells identified as arrested in the S phase by DNA ploidy (DAPI staining) was increased by the combination (Figure 6B, *right*). The arrest in the S phase coincided with an increased apoptosis induced by IM+SenB treatment (Figure 6A).

These results suggested that cells treated with IM+SenB could not undergo IM- induced cell cycle arrest and entered the S phase, apparently causing RS that contributed to S phase arrest and cell death. To investigate this hypothesis, we measured RS markers in IM- *vs* IM+SenB-treated K562 cells at 12 h (the beginning of cell cycle arrest) and 24 h (the time point with the increased S phase arrest and apoptosis). Figure 6C shows the increase of phosphorylated RPA32, a RS marker which binds single-strand DNA (ssDNA) and is hyperphosphorylated upon DNA damage or replication stress (58). Elevation of pRPA32 coincided with increased S345 phosphorylation of Chk1, a key kinase in the DNA damage response activated by ssDNA damage and stalling of replication forks. Interestingly, the DNA repair proteins BRCA1 and Rad51, which are recruited to ssDNA sites, were downregulated by IM and further downregulated by IM+SenB at 24 h, indicating inhibition of DNA repair by these treatments (Figure 6C).

## Discussion

Treatment of CML with IM remains one of the most efficient anticancer therapies to date. Despite its success, patients receiving BCR-ABLi must continue treatment to avoid relapse, due to persistence of resistant quiescent LSC (59). While BCR-ABL mutants are successfully treated with new generations of target inhibitors, non-mutational resistance, and in particular quiescence-driven resistance remains a challenge (60).

The Mediator kinases CDK8/19 regulate effects of a variety of exogenous stimuli. CDK8/19i have been established as anticancer drugs, active either alone or in combination with different therapies. In the present article, we demonstrated that CDK8/19i SenB and SNX631 sensitized K562 CML cells to various BCR-ABLi (Figures 2A-D). The most significant effect of this sensitization was the increase in the rate of apoptosis (Figure 1). It was previously reported that CDK8 is downregulated by dasatinib but not by IM, and that siRNA knockdown of *CDK8* sensitizes K562 cells to IM but not to dasatinib (61). In the present study, we observed no changes of CDK8 levels by 24 h treatment with IM (Figure S10), and we found that CDK8/19i sensitized K562 to all BCR-ABL antagonists including dasatinib (Figure 2). A likely cause for the different results obtained with *CDK8* siRNA and CDK8/19i is that the latter also inhibits CDK19, which exerts the same effects as CDK8 (29).

IM inflicted global changes in gene expression affecting several pathways, especially those controlling cell proliferation and interferon/STAT signaling (Figure 3C). While the combination had no dramatic effect on pathways most sensitive to IM alone (Figure 3C), the addition of CDK8/19i markedly shifted the effects of IM on gene expression, counteracting the downregulation but enhancing the upregulation of gene expression caused by the BCR- ABLi (Figures 3А-B). These results resemble the effects of CDK8/19i on castration-induced changes in gene expression in prostate cancers (44), reflecting both the positive and the negative effects of CDK8/19 on transcription (29).

Activation of pro-survival pathways contributes to tumor cell persistence. Importantly, in at least 25% of cases and up to 60% of resistance to BCR-ABLi stems from mechanisms independent of the structure of the *BCR-ABL* indicating BCR-ABL-independent mechanisms (60). The major pro-survival pathways activated in response to BCR-ABL inhibition are STAT3 and STAT1 (62). In turn, inhibition of these pathways can increase the potency of BCR-ABLi. We found that the combination of SenB and IM reduced the amounts of total STAT1 and S727-phosphorylated STAT1 and STAT3 (Figures 2B,D,E). JAK2/STAT3 is a major contributor to BCR-ABLi resistance, activated by autocrine signaling (10) and bone marrow-secreted cytokines(9). BCR-ABL dependent STAT3 S727 phosphorylation that augments STAT3 transcriptional activity contributes to resistance to BCR-ABLi (63). STAT3 inhibition can overcome resistance to BCR-ABLi and induce synthetic lethality in STAT3-dependent CML including LSCs (64). The effect on STAT3 phosphorylation could serve as one of the mechanisms of the combinatorial effect of BCR- ABLi and CDK8/19i.

However, the most dramatic effect of the addition of CDK8/19i to BCR-ABLi revealed in this study was the effect on the induction of quiescence. From the onset of the TKI era in CML treatment, it became clear that non-dividing cells are much less sensitive to these drugs than the proliferating counterparts. Subpopulations of quiescent LSCs survived the exposure to BCR-ABLi resulting in disease relapse (65). IM induced apoptosis primarily in S and G2/M phases (66), therefore investigational strategies focused on eliminating quiescent cells or inhibitor-induced G1 cell cycle arrest. Altered response of CML cells to IM via quiescence has been reported (15,65). Quiescence has been implicated in the delay of full clinical remission (15,16) and in the disease relapse (17).

Our major mechanistic finding is the identification of IM-induced G1 cell cycle arrest as a critical factor in CML cell sensitization by CDK8/19i. IM treatment of K562 cells led to G1 accumulation (Figure 6A, *top left*) and elevation of CKIs p27^Kip1^ and p18^INK4c^ (Figures 4B-C). Of note, p27^Kip1^ and p57^Kip2^ were inducible within the initial hours of treatment with IM (24). Furthermore, CKIs have been mechanistically related to LSC persistence (19,67). Strikingly, combining IM with a CDK8/19i released the cells from G1 arrest, prevented p27^Kip1^ accumulation (Figures 4A, *right*, 4B, 4C, *left,* 5С) and enhanced the pro-proliferation driver c-Myc mRNA and protein levels (Figures 4A, *left* and 4C, *left*) and c-Myc-dependent transcripts (Figure 3C). This observation was supported by the results that the inducible p27^Kip1^ expression in K562 cells increased the G1 fraction and attenuated IM-induced apoptosis. Although both effects were reversible by combining IM with SenB (Figure 5), protection from p27^Kip1^ induction was not completely abrogated by CDK8/19i. This points to an alternative explanation – that downregulation of p27^Kip1^ is correlated with an increase of replication rate and could be the consequence of cells entering replication, rather than its cause, suggesting that CDK8/19i counteracts cell cycle arrest through alternative mechanisms. The addition of SenB to IM elevated the fraction of cells entering replication (Figure 6B, *left*, S9), but the ensuing replication was halted and RS markers such as pRPA32/RPA2 and pChk1 were induced, concurrently with the onset of apoptosis.

RS in BCR-ABL-positive cells is increased without treatment (68) and contributes to IM resistance (69). On the other hand, CML cells appear to be sensitive to the combination of IM and DNA damaging treatments such as inhibition of NEDD8 activating enzyme (NAE1) (70) and Rad51 (71). Similarly to our combinations, NAE1 inhibition increased the fraction of EdU-negative cells in S phase, and the inhibitors of NAE1 and Rad51 in combinations with IM significantly increased apoptosis. Also, RS in response to hydroxyurea sensitized CML cells to IM (72). Interestingly, we found that DNA repair proteins Rad51 and active (phosphorylated) BRCA1 were downregulated by the combination of IM+SenB (Figure 6C).

Supporting the role of cell cycle regulation in the sensitivity to IM and the sensitization by CDK8/19i, IM influenced neither cell cycle progression nor CKI and c-Myc protein expression (Figures 4C, *right*, 6A, *right*) in BCR-ABL-positive KU812 cells. Rather, these cells readily underwent IM-induced death in a CDK8/19-independent manner.

Similarly to our results, Nakamura et al. demonstrated that CDK8/19i-induced premature G1-S transition can promote DNA damage and RS (48). On the other hand, Xu et al. reported that CDK8 acted as a negative regulator of p27^Kip1^ stability but did not affect its mRNA level in breast cancer cells (73). Results that we obtained in CML cells differ from these observations, suggesting that the effect of CDK8 on p27^Kip1^ is cell type-specific.

Altogether, we demonstrated that pharmacological inhibition of CDK8/19 promotes apoptosis in CML cells treated with BCR-ABLi by attenuating BCR-ABLi-induced cell cycle arrest thereby preventing the escape from apoptosis (Figure 6D). The mechanism of such sensitization demonstrates an example of a general principle, where increase in the rate of cell proliferation by CDK8/19i in combination with BCR-ABLi elicits a pro-apoptotic response through inhibition of pro-survival properties of quiescence (58). Our results support the perspective of combining the inhibitors of BCR-ABL and CDK8/19 for the treatment of CML.

## Author Contributions

A.I.K., M.Y., and V.T. conceived and conducted the study, performed the majority of experiments, analyzed data, prepared figures and graphs, and wrote the original draft. A.B. and M.Z. obtained the K562p27^tet-on^ subline. C-U.L. and J.N. performed additional flow cytometry experiments. N.V. prepared cDNA libraries for RNASeq, A.M.K., E.V., A.T., and J.L. analyzed RNASeq data. E.I. and Y.A. performed additional immunoblotting experiments. I.R., E.B., M.C., and A.S. discussed the project, reviewed and edited the manuscript. The text was read and approved by all authors.

## Conflict of Interest

I.R. is Founder and President, M.C. is an employee, and E.B. is a consultant of Senex Biotechnology, Inc. Other authors declare no conflict of interest.

## Acknowledgements

We thank the Center for Precision Genome Editing and Genetic Technologies for Biomedicine, Institute of Gene Biology, Russian Academy of Sciences, for the research facility support. We thank Dr. Artem Velichko for the antibodies to RPA32/RPA2, pRPA32/RPA2 S4/S8 and Rad51. The work was supported in part by the Microscopy and Flow Cytometry Core (MFCC) of the Center for Targeted Therapeutics at the University of South Carolina, Columbia, SC (2019-2021).

## Funding

This work was supported by Megagrant (Agreement №14.W03.31.0020 between the Ministry of Science and Education of the Russian Federation and Institute of Gene Biology, Russian Academy of Sciences).

## List of abbreviations

BCR-ABLi: BCR-ABL inhibitor(s);
CDK: cyclin-dependent kinase;
CDK8/19i: CDK8/19 inhibitor(s);
CKI: cyclin-dependent kinase inhibitor;
CML: chronic myelogenous leukemia
Das: dasatinib;
DEG(s): differentially expressed gene(s);
Dox: doxycycline;
FC: fold change;
FDR: false discovery rate;
IM: imatinib mesylate;
LSC: leukemia stem cells
Nilo: nilotinib;
PARP: poly (ADP-ribose) polymerase
RS: replication stress;
SenB: Senexin B.

## Notes

### Summary of Updates

In the current version there are some corrections and updates as compared with the manuscript v.1. Figures 5 and 6 and the related text in Results and Discussion were revised and supplemented with a new data. The role of CDK8/19 inhibition in cell cycle regulation of CML has been demonstrated more distinctly. Missed biological repeats and statistical processing of data were added where needed. Section Competing Interests was updated.

